# Human muscle stem cells are refractory to aging

**DOI:** 10.1101/2020.08.07.238477

**Authors:** James S. Novak, Davi A.G. Mázala, Marie Nearing, Nayab F. Habib, Tessa Dickson, Olga B. Ioffe, Brent T. Harris, Marie N. Fidelia-Lambert, Christopher T. Rossi, D. Ashely Hill, Kathryn R. Wagner, Eric P. Hoffman, Terence A. Partridge

## Abstract

Age-related loss of muscle mass and strength is widely attributed to limitation in the capacity of muscle resident satellite cells to perform their myogenic function. This idea contains two notions that have not been comprehensively evaluated by experiment. First, it entails the idea that we damage and lose substantial amounts of muscle in the course of our normal daily activities. Second, it suggests that mechanisms of muscle repair are in some way exhausted, thus limiting muscle regeneration. A third option is that the aged environment becomes inimical to the conduct of muscle regeneration. In the present study we used our established model of human muscle xenografting to test whether muscle samples taken from cadavers, of a range of ages, maintained their myogenic potential after being transplanted into immunodeficient mice. We find no measurable difference in regeneration across the range of ages investigated up to 78 years of age. Moreover, we report that satellite cells maintained their myogenic capacity even when muscles were grafted 11 days postmortem in our model. We conclude that the loss of muscle mass with increasing age is not attributable to any intrinsic loss of myogenicity and is most likely a reflection of progressive and detrimental changes in the muscle micro-environment such as to disfavor the myogenic function of these cells.

## Introduction

Sarcopenia, the age-related loss of skeletal muscle, is the most common type of muscle atrophy in humans leading to significant impairments in function and strength ^1^. In humans, sarcopenia typically begins to intrude around the 4^th^ decade of life at a rate of roughly 1% muscle loss annually, progressively amounting to a loss of 30–50% skeletal muscle mass by 80 years of age ^2,3^. This process is heavily influenced by extrinsic factors stemming from the inflammatory, fibrogenic and adipogenic cells that constitute the muscle interstitial microenvironment ^4,5^. With age, the propagation of pro-inflammatory cytokine signaling and infiltration of fibrotic and adipogenic tissue within the muscle, together, have been shown to exert detrimental impacts on muscle satellite cell (MuSC) quiescence and regenerative capacity ^4–9^. Sarcopenia is intimately associated with many age-related pathologies, including frailty, Alzheimer’s disease, type 2 diabetes, and others ^10–12^.

Although a number of investigations of the basis of sarcopenia have demonstrated a major extrinsic component to myogenic competence in aged individuals ^13–18^, other studies have implicated intrinsic defects in the behavior of aged MuSCs that play an important part in the failure to maintain muscle mass, MuSC senescence and regenerative capacity with aging ^16–18^. These postulated cell-intrinsic deficits that impact myogenic competence with aging may arise from several factors, including depletion of the aged MuSCs due to increased asymmetric divisions within the heterogenous population ^19,20^, and alterations in cell-cycle/differentiation signaling cascades, such as the p38/MAPK axis that negatively regulate MuSC self-renewal through inhibition of *Pax7* and activation of *Myod1* ^21,22^. However, much of this work has been conducted in rodent models where proliferative persistence despite aging has been associated with long telomeres ^23–26^.

Certainly, amongst mammalian tissues, skeletal muscle is highly sensitive to the aging process and represents an impressive model in which to dissect the intrinsic and extrinsic components of age-associated changes in tissue homeostasis and function ^27^. Previously, we have shown that skeletal muscle from Facioscapulohumeral muscular dystrophy (FSHD) patients and unaffected relatives can regenerate effectively within the anterior tibial compartment of immunodeficient host mice ^28^. These human donor grafts regenerate into muscle fibers containing >90% human myonuclei, together with a micro-vascular bed of dual mouse/human origin and innervation from the host anterior tibial nerves ^28^. This system allowed us to successfully transplant muscle harvested postmortem from human cadavers, that was indistinguishable from that of grafts of living donors ^28^. This is in accord with a report showing the rescue of viable and vigorously myogenic MuSCs from muscle several days post mortem ^29^. Although it has been suggested that myogenic cells remain viable within a muscle for around 2 weeks postmortem ^29^, the extent to which this population maintains full regenerative potential across an increasing postmortem interval (PMI) after death has yet to be investigated, where survival of other cell types, such as fibro-adipogenic cells are increasingly understood as playing important accessory roles ^30–34^.

Here we report further development of the xenograft model to permit an *in vivo* evaluation of the myogenic reserves of human skeletal muscle collected across a range of ages and postmortem intervals. Here, we hypothesized that human muscle derived from either young, middle-age or old human cadavers would graft and regenerate equally effectively following the same time-courses of proliferation, differentiation and myofiber regeneration. Additionally, we hypothesized that MuSCs would maintain their myogenic potential after death at increasing postmortem intervals. This revealed no measurable difference in human muscle regeneration across the range of ages investigated up to 78 years of age, and further, report that MuSCs maintained their myogenic capacity even when muscles were grafted up to 11 days postmortem. We conclude that the loss of muscle mass with increasing age is not attributable to any intrinsic loss of myogenicity of the MuSCs and is most likely a result of persistent detrimental alterations in the multi-cellular muscle environment. This study has provided a stringent examination of the extent to which the loss of muscle mass in aged and cachexic individuals reflects intrinsic defects in the proliferative or differentiative functions of the MuSC population present in the context of the aged muscle.

## Results

### Xenograft model of human muscle satellite cell-mediated regeneration

In this study, the primary objectives were to comprehensively evaluate the influence of human age and postmortem interval on the regenerative capacity of human MuSCs following transplantation in a xenograft mouse model. Human postmortem muscle samples were procured with approval by the Children’s National Hospital Institutional Review Board in collaboration with regional medical centers. Samples were described only by age, gender, race, and date/time of death and autopsy. Exclusion criteria for human samples included only patients with blood-borne infectious diseases (i.e. HIV, Hepatitis) or neuromuscular diseases. Xenograft transplantation was performed in accordance with work with our previous collaborators ^28^. In this study, we obtained a total of 13 human muscle samples both males and females ranging from 28 to 91 years of age (**Table 1**). Human muscle samples were predominantly isolated from the psoas muscle, but also included samples of abdominal and rectus femoris muscles (**Table 1**). In brief, human donor muscle samples were dissected at autopsy under sterile conditions and transported to Children’s National Hospital. They were trimmed into multiple ~8 mm length strips, approximating the size of the mouse tibialis anterior muscle, and transplanted into the anterior compartment of NOD.Cg-*Rag1*^*tm1Mom*^ *Il2rg*^*tm1Wjl*^/SzJ (NRG) immunodeficient mice ^28,35^. A single human muscle strip was then sutured to the distal and proximal tendons of the mouse extensor digitorum longus muscle and the graft site was closed with surgical glue and wound clips.

**Table 1.** Human muscle samples and xenograft study results based on donor age and postmortem interval. Description of the age, sex, race, muscle sample, postmortem interval (PMI), and rate of success for all human cadaver samples investigated in this xenograft study of human skeletal muscle regeneration.

| Age | Sex | Race | Muscle | PMI | Success Rate (n per PMI) |
| --- | --- | --- | --- | --- | --- |
| 28y | Female | African American | Psoas | 4d, 8d, 12d | 0/8 (4d), n/a (8d), n/a (12d) |
| 36y | Male | Caucasian | Psoas | 5d; 7d | 2/8 (5d), 1/9 (7d) |
| 41y | Male | African American | Psoas | 3d, 5d | 0/5 (3d), 0/1 (5d) |
| 54y | Male | Caucasian | Psoas | 2d, 4d | 1/2 (2d), 1/3 (4d) |
| 58y | Male | African American | Psoas | 3d, 6d | 2/2 (3d), 4/4 (6d) |
| 66y | Female | *Not specified* | Abdominal | 4d, 11d | 8/10 (4d), 7/11 (11d) |
| 68y | Male | *Not specified* | Psoas | 5d, 7d | 0/5 (5d), 0/4 (7d) |
| 70y | Female | Caucasian | Psoas | 3d, 5d | 0/2 (3d), 0/1 (5d) |
| 70y | Female | *Not specified* | Psoas | 5d, 6d, 7d | 0/1 (5d), 0/3 (6d), 0/2 (7d) |
| 75y | Male | *Not specified* | Abdominal | 2d, 3d, 6d | 4/4 (2d), 2/2 (3d), 3/3 (6d) |
| 77y | Male | *Not specified* | Abdominal | 2d, 6d | 2/3 (2d), 0/2 (6d) |
| 78y | Male | African American | Rectus Femoris | 11d | 4/6 (11d) |
| 91y | Male | *Not specified* | Psoas | 2d, 9d, 16d | 0/5 (2d), 0/5 (9d), n/a (16d) |

Regeneration of the human muscle was examined 3- or 6-weeks post-engraftment to investigate efficiency of muscle regeneration with donor age and the postmortem interval between death and xenograft transplantation. Overall, successful regeneration of the human muscle was found with 7 of the obtained human muscle samples in this study, at postmortem intervals between 2 and 11 days (**Table 1**). Successful regeneration of the human muscle samples was confirmed by immunostaining for human membrane protein spectrin and nuclear protein lamin A/C (**Supplementary figure 1a**). While not all muscle grafts successfully regenerated to form myofibers within the host, in many of these samples we did find evidence of human spectrin and lamin A/C that localized within areas of connective tissue adjacent to regenerated host muscle (**Supplementary figure 1b**). In some cases, failure to regenerate was associated with infection attributable to contamination during isolation of the donor muscle or during the engraftment procedures. Assessment of regenerative success (i.e. number of human myofibers and area) was complicated by variation along the length of the muscle (**Supplementary figure 1c**).

### Grafted human muscle satellite cells maintain myogenic capacity irrespective of age

In order to first investigate the influence of age-related sarcopenia on the regenerative capacity of human MuSCs, we performed xenograft transplantation of postmortem human muscle samples from individuals ranging from 28 to 91 years of age. Analysis of xenografts at 3- or 6-weeks following engraftment indicated successful regeneration across the range of donor age, plentiful regenerated myofibers of human origin being identified at sites of grafts from donors of 36, 54, 58, 66, 75, and 78 years of age (**Table 1**). Further, the most robustly regenerative human muscle samples, consistenty yielded both the the greatest numbers of regenerated human myofibers and the largest of contiguous areas of regenerated fibers (**Figure 1a-h; Supplementary Figure 2**) at 2 or more graft sites. These results were found across the range of ages (58-, 66-, 75- and 78-year old) of human samples. Thus our xenograft model revealed no compromised myogenic capacity of human MuSCs with age (**Figure 1a-d; Supplementary Figure 2**). Across the range of ages tested we also noted no differences in initial rate of regeneration or subsequent maturation of these human myofibers over the first 3 weeks post-engraftment (**Figure 1a-h**). In fact, one of our most successful xenografts were derived from our 78-year old human muscle sample, producing an average of 537 human myofibers across multiple successful transplants, and a study-wide maximum of 1019 regenerated human myofibers (**Figures 2 and 3**). Observations at 6-weeks post-engraftment revealed mature myofiber clusters as identified by a predominance of human myonuclei located in a juxta-sarcolemmal position (**Figure 3a, c**), together with numerous small-diameter myofibers with centrally-placed myonuclei suggesting a more recent bout of regeneration perhaps involving fiber branching (**Figure 3a, d**). This indicates that regeneration continues for at least 6 weeks beyond the initial period of re-growth over the first 3 weeks following engraftment. The patches of small centrally nucleated fibers seen at 6 weeks were localized to zones around the blocks of mature fibers and were intermixed with inflammatory or interstitial connective tissue associated with surgery. Despite the lack of evidence of regenerated human myofibers in our 28- and 91-year old cohorts that correspond to minimum/maximum ages within our associated study (**Table 1**), our data does not suggest any deficit in regenerative capacity of human MuSCs associated with increasing human muscle age in this xenograft model.

**Figure 1.**
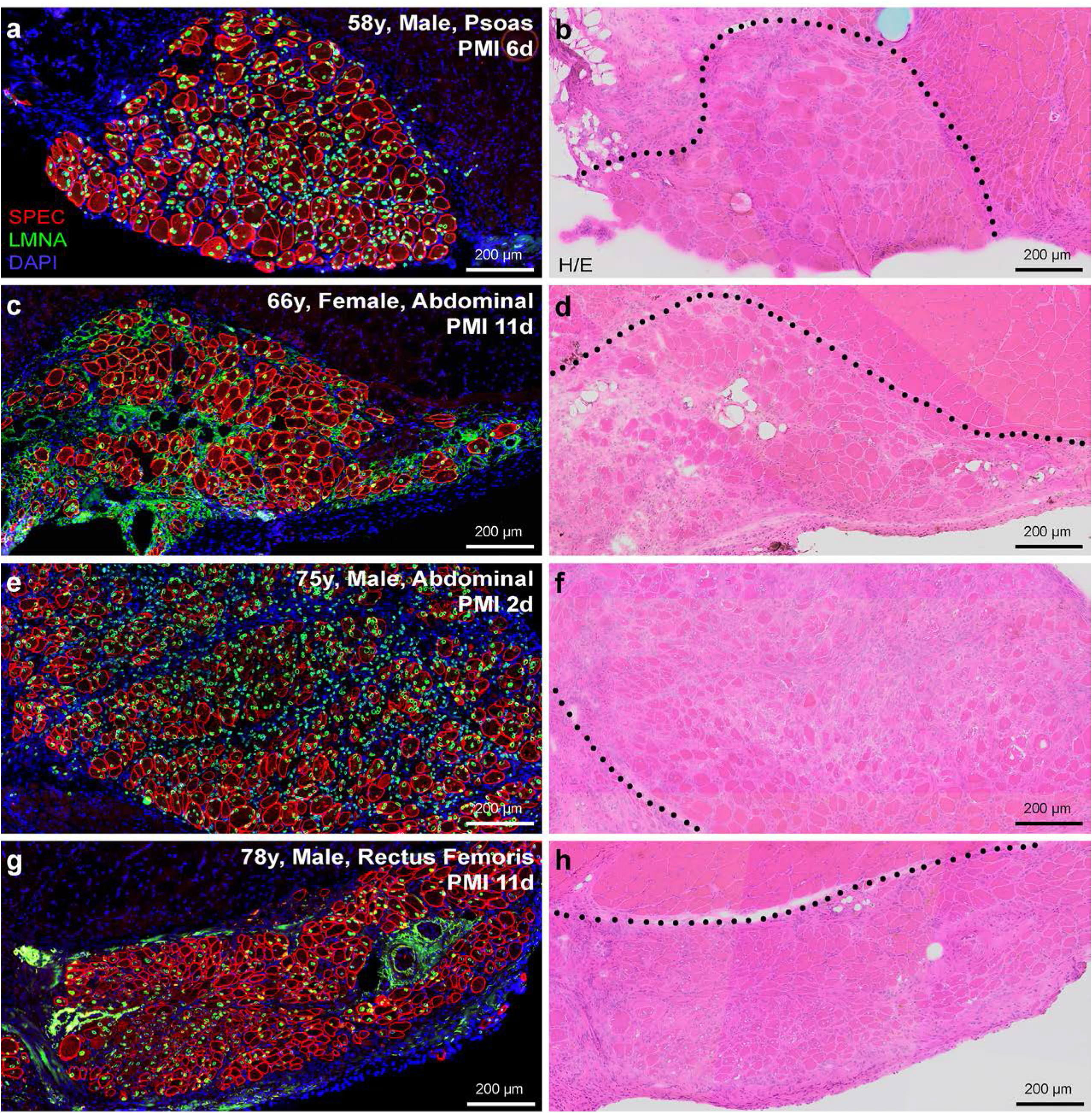
Myogenic capacity of human MuSCs is uncompromised despite aging. **a-h)** Human skeletal muscle regeneration following postmortem muscle tissue engraftment from human cadavers of various ages and postmortem intervals. Immunostaining for human-specific spectrin (SPEC, red) and lamin A/C (LMNA, green) proteins identify regions regenerated from dormant human MuSCs within the grafted tissue, while DAPI (blue) mark the entire muscle tissue section for reference (**a, c, e, g**). Human muscle regeneration shown here from transplanted human cadaver muscle at the indicated postmortem interval (PMI); 58-year old (**a, b**; male, psoas), 66-year old (**c, d**; female, abdominal), 75-year old (**e, f**; male, abdominal), and 78-year old (**g, h**; male, rectus femoris). Histology of adjacent serial sections performed by H/E staining shows regrowth of both human and mouse muscle tissue (delineated by black dotted line) harvested from anterior tibial compartment 3-weeks post-engraftment (**b, d, f, h**). Scale bars represent 200 μm (**a-h**).

**Figure 2.**
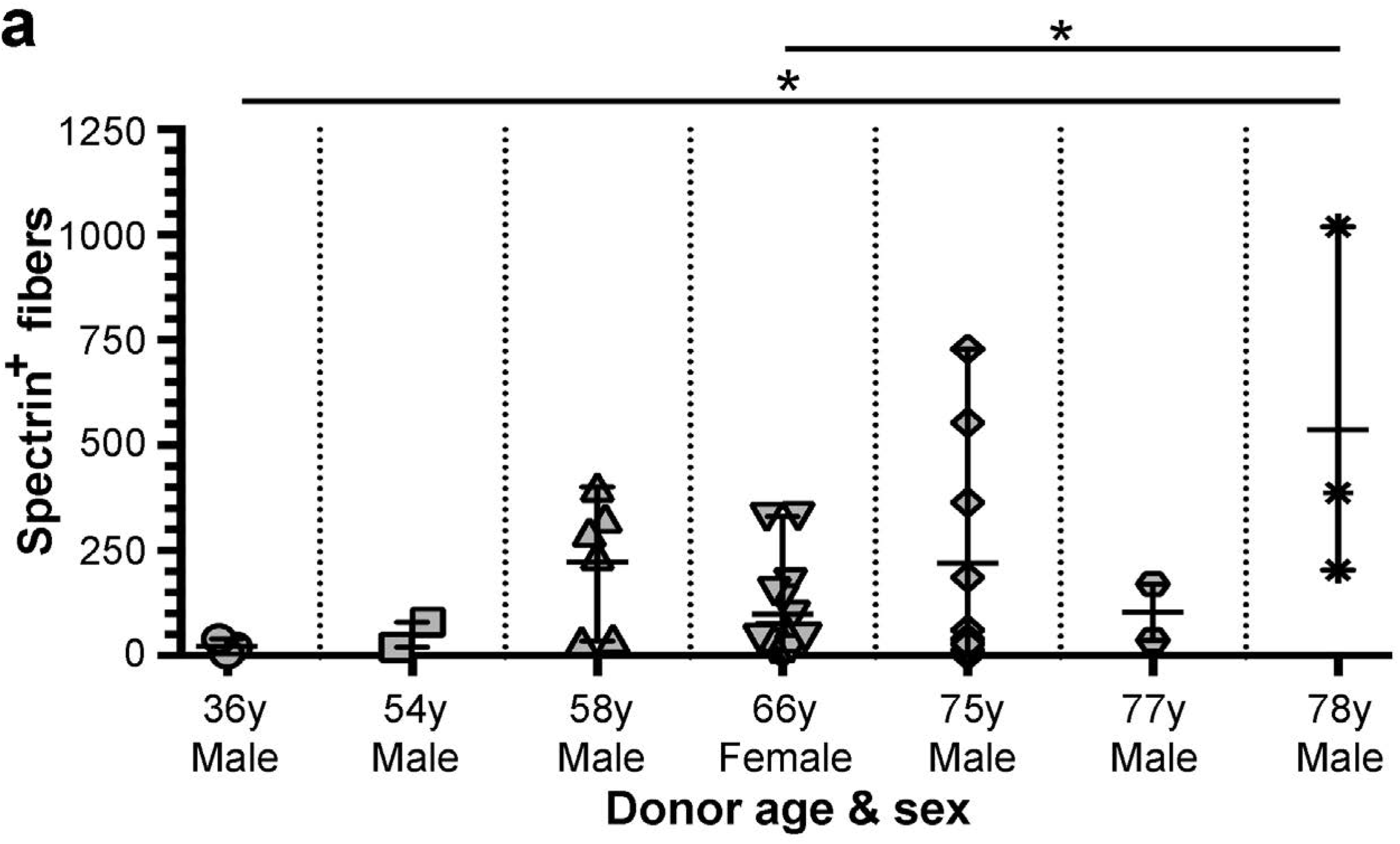
Assessment of human muscle regeneration by donor age and sex. **a)** Regenerated human skeletal muscle fibers were quantified based on proper localization of human-specific spectrin and lamin A/C proteins. Data represented as scatter-plot with mean and standard deviation (S.D.). for all successfully-regenerated cadaver samples as indicated in Table 1. Statistical analyses performed by one-way ANOVA with multiple comparisons; \**P* < 0.05.

**Figure 3.**
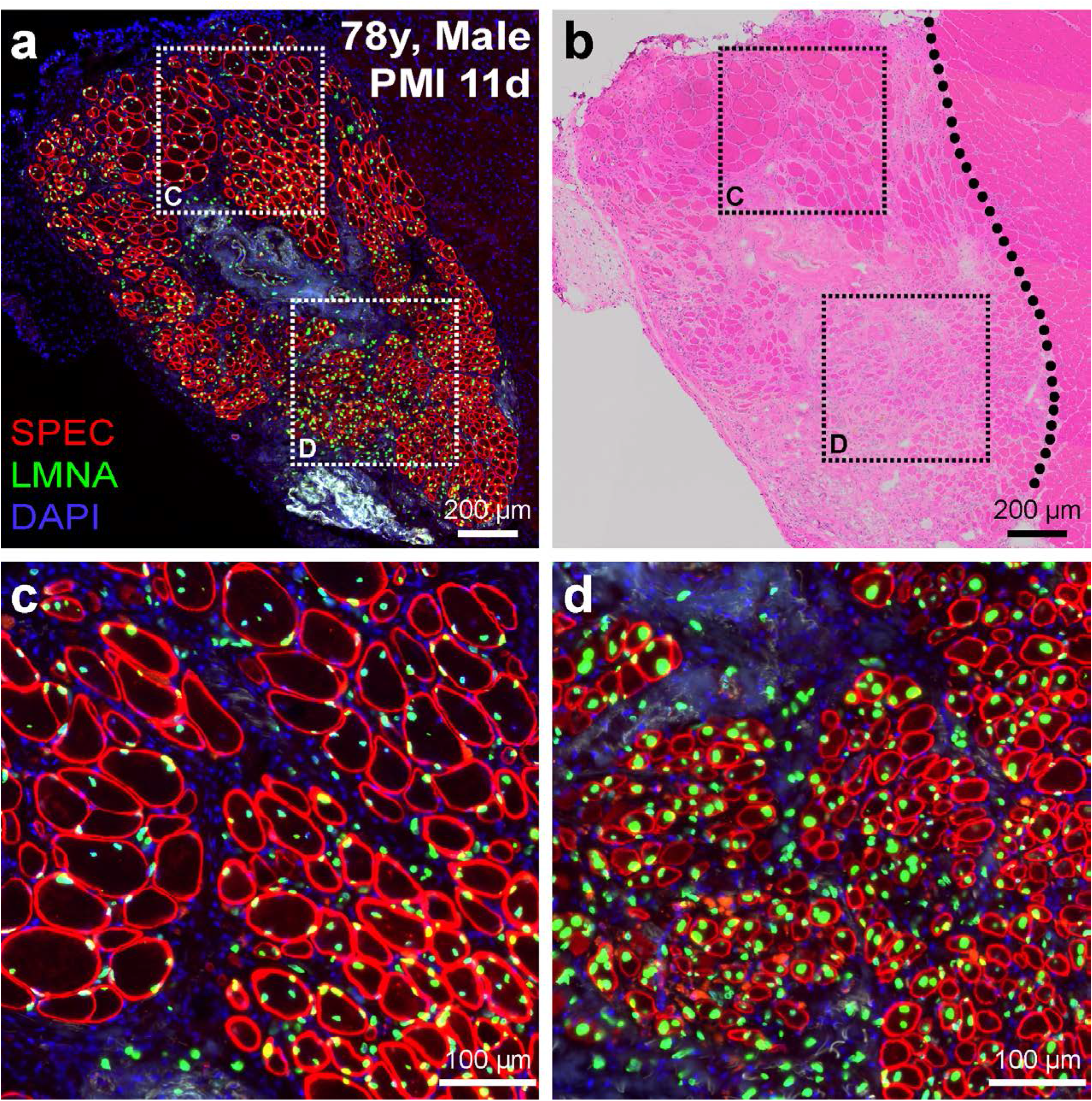
Efficient human muscle regeneration in xenograft of 78-year old cadaver muscle sample transplanted 11 days postmortem. **a, b)** Human skeletal muscle regeneration following postmortem (PMI, 11 d) muscle tissue engraftment from 78-year old human cadaver 6-weeks post-engraftment. Immunostaining for human-specific spectrin (SPEC, red) and lamin A/C (LMNA, green) proteins (**a**) indicate the extent of regeneration in this ‘high-responder’ xenograft, while H/E staining (**b**) shows the boundary of human muscle regeneration adjacent to the regeneration mouse muscle tissue (delineated by black dotted line). **c, d)** Inlays represent areas of mature fibers marked predominantly with peripheral nuclei labeled with human lamin A/C (**c**) or newly-regenerated, centrally-nucleated fibers as indicated by the position of lamin A/C-labeled myonuclei (**d**). Scale bars represent 200 μm (**a, b**) or 100 μm (**c, d**).

Analysis of engraftment efficacy by sex or race also did not reveal any major differences in the survival or regenerative capacity of human MuSCs. Of the 13 human muscle samples collected, 4 were from female cadavers while 9 were from male cadavers. While we failed to find evidence of regeneration in 3 of 4 female samples, the 66-year old female sample compared well with the successful male grafts in terms of its regenerative capacity (**Table 1; Figure 2**). Taken together with the relatively short postmortem interval tested for all female samples (PMIs ranging 2-5 days), it would suggest potential issues with the health status of the muscle sample (contamination or compounding illness) or surgical procedure rather than any link to sex. Of the 9 male muscle samples evaluated in our study, we failed to find evidence of regeneration in 3 of the xenograft surgeries performed; however, we found that all three 70-80-year old male samples regenerated well in our xenograft model (**Table 1; Figure 2**). Stratification of the data by race also did not point towards any apparent link with regenerative capacity. In our study, we evaluated 4 African American muscle samples (2 of which regenerated) and 3 Caucasian muscle samples (2 of which regenerated), while 6 were not specified in terms of donor race (**Table 1; Figure 2**). The number of samples received limits our evaluation in terms of this demographic; however, we see no indication of any major differences based on sex or race in this study.

### Myogenic capacity is preserved with increasing postmortem intervals

MuSCs can adopt a quiescent state and retain their regenerative capacity postmortem ^29^. This intriguing cellular adaptation within the seemingly hostile, necrotic environment of a postmortem cadaver muscle occurs under conditions of anoxia and nutrient deprivation ^29^. Furthermore, the postmortem isolation of MuSCs at intervals from both murine and human muscle demonstrated retention of their proliferative and regenerative capacities ^29^ for several days. Here, another major primary objective was to investigate retention of regenerative capacity of human MuSCs maintained under anoxic conditions for extended periods. Cadaver muscle samples were received at 2-11 days post mortem and stored in Dulbecco’s Modified Eagle’s Medium (DMEM) at 4±C until grafting – the total time being defined as our postmortem interval. In the 5 of our 7 human cadaver muscle samples that successfully regenerated *in vivo*, we grafted at a number of PMIs whenever possible, depending on the size of the sample and the availability of suitable recipient mice. In all human muscle samples that successfully regenerated *in vivo*, we found no apparent decline in the regenerative capacity of human MuSCs with extended PMIs (**Figure 4a-g**). Most notably, longitudinal assessments of the 66-year old human muscle samples at PMIs of 4 and 11 days indicated preserved myogenic integrity with regard to the amount of muscle regeneration we observed in this model (**Figure 4a, b, g**). Further, assessment of the 78-year old sample at a single PMI of 11 days yielded our most extensive regenerates of human muscle, indicating no loss in postmortem regenerative capacity with both advanced age and prolonged PMI (**Figure 4e, f, g**). In both these cases, PMIs out to 11 days were found to regenerate efficiently *in vivo* (**Figure 4g**). In the case of the 75-year old sample, we found relatively low numbers of regenerated human fibers in our third engraftment attempt at a PMI of 6 days but the low n-value leaves us unable to evaluate the possibility of sampling error (**Figure 4c, d, g**). Overall, our data stratified by PMI for all successful human samples indicate no conspicuous loss of MuSC regenerative capacity out to nearly 2-weeks postmortem, and suggests the use of postmortem muscle tissue and its resident MuSCs for future experimental and translational regenerative studies.

**Figure 4.**
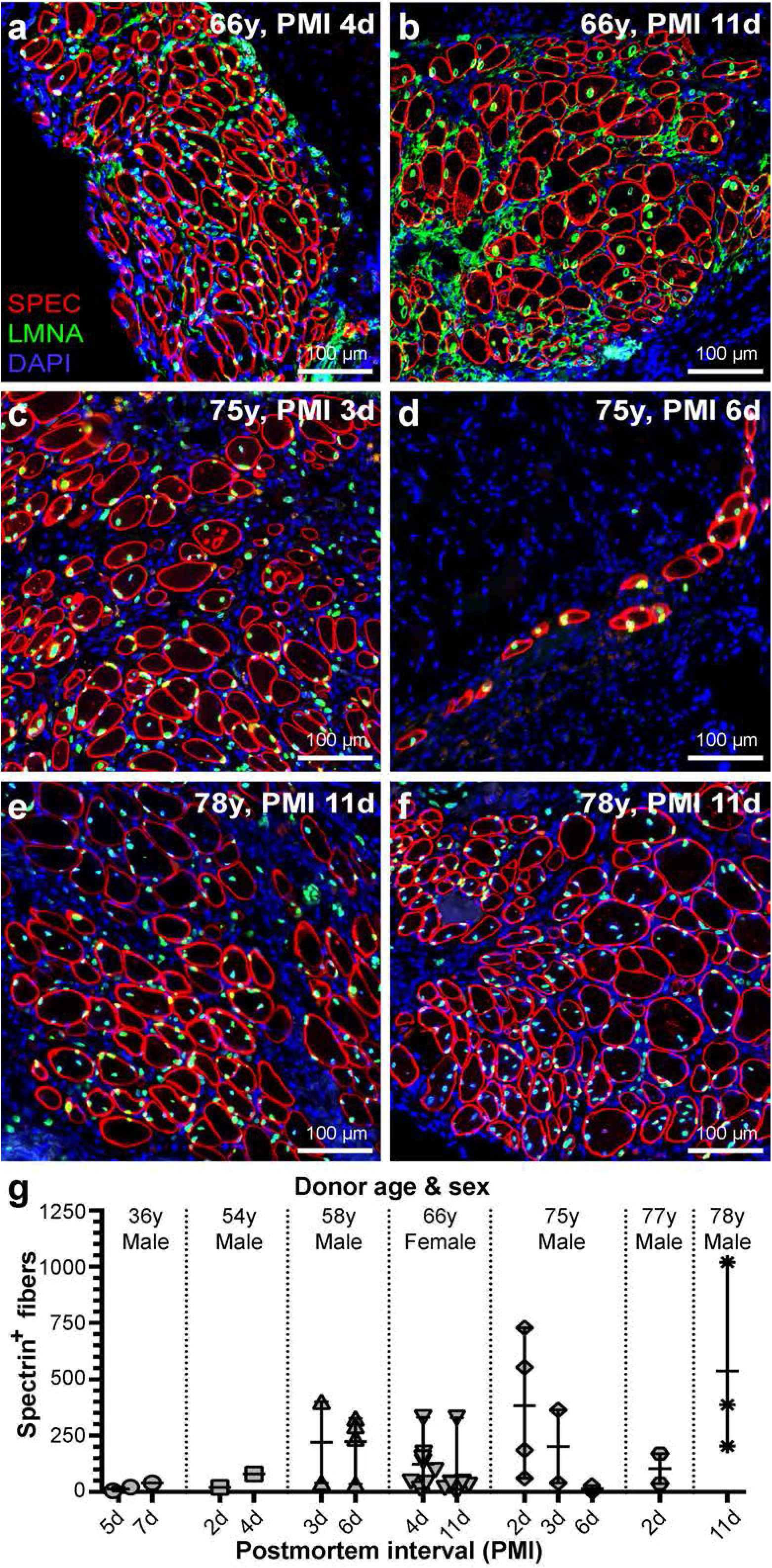
Resiliency of grafted human MuSCs after prolonged postmortem dormancy. **a-g)** Assessment of human muscle regeneration at escalating postmortem intervals (PMI). Comparisons at two PMIs for the 66-year old sample (**a, b**; PMI 4 d and 11 d) and 75-year old sample (**c, d**; PMI 3 d and 6 d) are shown. Successful regeneration after 11 days postmortem is demonstrated in two xenografts for the 78-year old sample indicating resiliency of human MuSCs despite prolonged PMI when maintained under anoxic conditions (**e, f**). Representative images indicate no apparent decline in MuSC survival or regenerative capacity out to 11 days postmortem (**b, e, f, g**). **g)** Regenerated human skeletal muscle fibers were quantified represented as scatter-plot with mean and S.D. for all successfully-regenerated cadaver samples stratified by PMI as indicated in Table 1. Statistical analyses were performed (when permitted by n-values) by non-parametric, Kolmogorov-Smirnov test to assess differences between PMIs for a given sample.

### Regenerated human muscle maintained growth and fiber size across all ages

Although we found no evidence of myogenic deficit with aging, we looked for deficits in myofiber growth between the 3- and 6-week post-engraftment cohorts. To assess this, we measured the minimal Feret diameter of all human fibers after 3- and 6-weeks post-engraftment and stratified the data by age in decades (i.e. 50-, 60-, and 70-year old samples). By using human-specific spectrin membrane immunostaining, we were able to differentiate muscle fibers derived from human MuSCs versus those derived from mouse MuSCs. While the size distribution of fibers from each cohort is strikingly similar following 3 weeks of growth post-engraftment, the distribution following 6 weeks demonstrates an increase in small-diameter fibers that may be attributable to a second wave of regeneration, as well as an increase in size of the more mature myofibers when compared to the data collected at 3-week post-engraftment (**Figure 5a, b**). The median minimal Feret diameter was very similar at 3 weeks post-engraftment in all groups (~20 μm), and overall 60% of the total fibers were less than 20 μm in diameter while 40% were greater than 20 μm. In contrast, at 6 weeks post-engraftment, while the combined median minimal Feret diameter of all groups was also ~20 μm (55% of the total fibers were less than 20 μm in diameter while 45% were greater than 20 μm), the median for each group ranged from 13 to 20 μm. The minimal Feret diameter from both the 50- and 60-year old cohorts was just slightly smaller on average at 6 weeks vs. 3 weeks post-engraftment due to increased presence of clustered small-diameter, newly-regenerated human fibers less than 15 μm in size (**Figure 5c**). In contrast, the average minimal Feret diameter of the 70-year old cohort was greater at 6 weeks vs. 3 weeks post-engraftment; however, we similarly observed clustered populations of both mature and small-diameter, centrally-nucleated fibers in this cohort similar to the other cohorts (**Figure 5b, c**).

**Figure 5.**
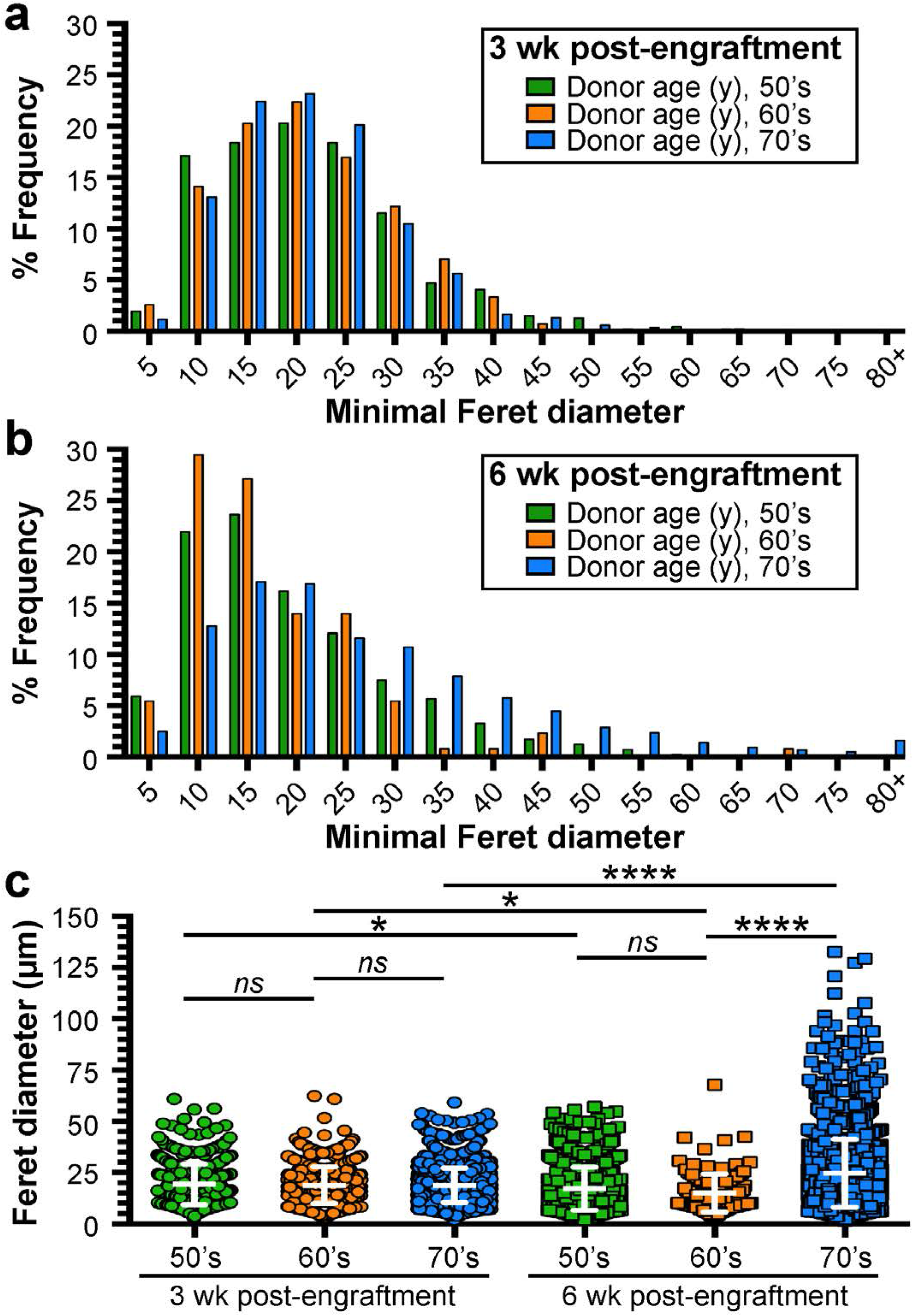
Analysis of muscle fiber size post-engraftment. Data was stratified by age in decades (i.e. 50-, 60-, and 70-year old samples). **a**) Minimal feret diameter quantified from samples stratified by age bracket harvested 3 weeks post-engraftment (50’s, n = 2 samples and 468 fibers; 60’s, n = 1 sample and 815 fibers; 70’s, n = 3 samples and 1884 fibers). **b**) Minimal feret diameter quantified from samples stratified by age bracket harvested 6 weeks post-engraftment (50’s, n = 2 samples and 829 fibers; 60’s, n = 1 sample and 130 fibers; 70’s, n = 3 samples and 2167 fibers). **c**) Scatter-plot showing mean with S.D. at both 3-weeks and 6-weeks post-engraftment for all cohorts. Statistical analyses performed by one-way ANOVA with multiple comparisons; *ns, not significant,* \**P* < 0.05; \*\*\*\**P* < 0.0001.

### Human myogenic cells migrate and regenerate beyond the initial graft site

Intriguingly, microscopic analysis of the regenerated human muscle at both 3- and 6-weeks post-engraftment indicate the persistence of human mononuclear cells (lamin A/C-positive) within the interstitial space between the regenerated human myofibers (**Figure 6a-d**; *solid arrows*). These human cells were localized strictly between zones of human and adjacent mouse muscle (**Figure 6a-d**). This may indicate the survival and expansion of various cell types within the regenerated human muscle that promote MuSC-mediated regeneration ^4,36^. Additionally, with prolonged growth after engraftment (i.e., 6 weeks after transplantation), we noted a reduction in the interstitial space between human fibers and the overall presence of these interstitial mononuclear cells (**Figure 6a-f**), as the inflammatory and fibrotic processes resolved with time. Further, in grafts examined 6-weeks post-engraftment, we observed the populations of human interstitial cells beyond the borders of human myofibers identified by human lamin A/C (*defined by dashed circular region*), as well as some low-intensity spectrin staining in what appear to be small-diameter fibers forming within this same region amongst the human lamin A/C-labeled mononuclear cells (**Figure 6c, d**; *indicated by ‘+’ signs*). This data may indicate continuing expansion of MuSCs and myofiber regeneration, even from our oldest human samples, for at least 6 weeks after transplantation.

**Figure 6.**
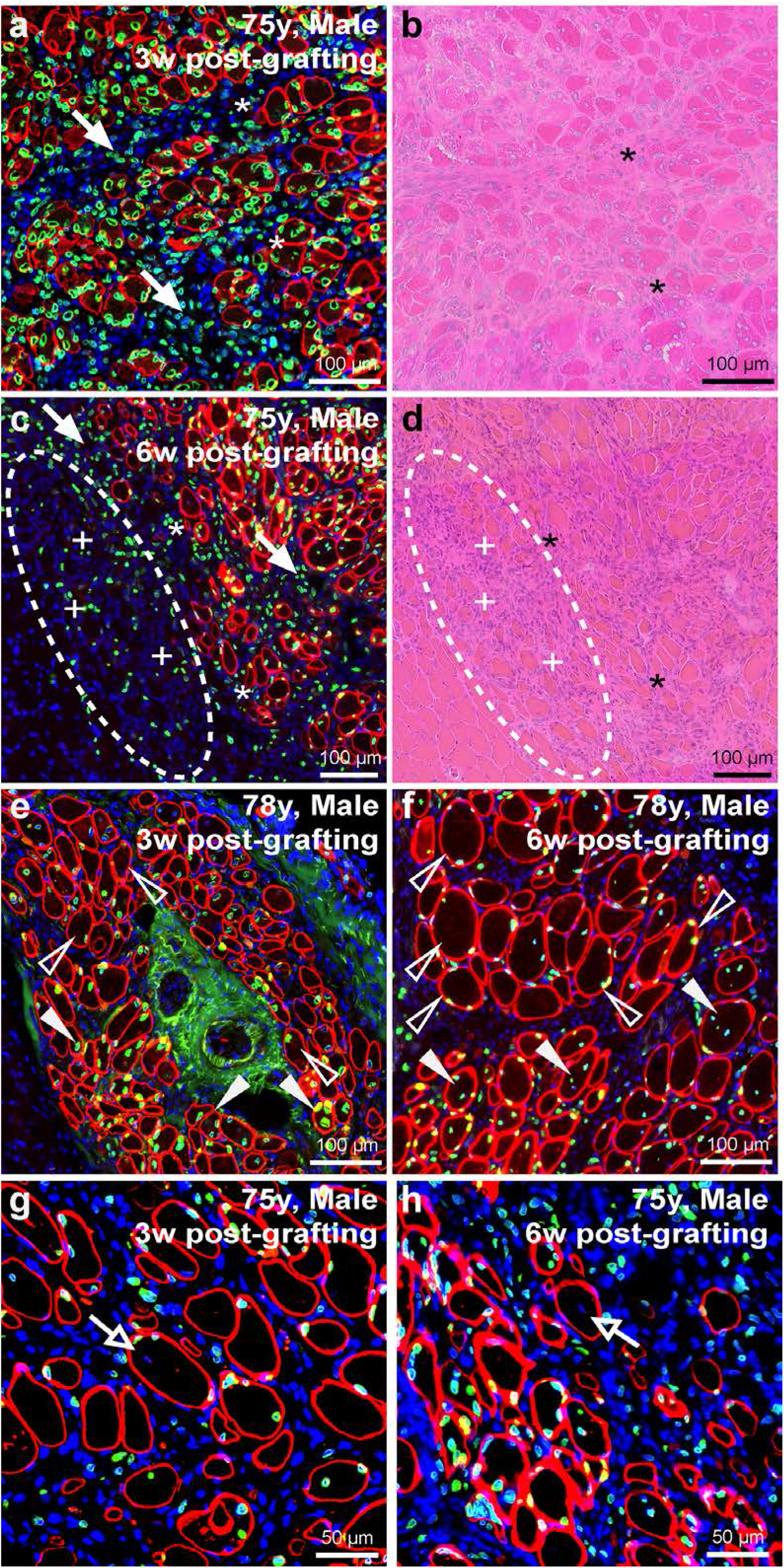
Myonuclear localization and migration post-engraftment. **a-d)** Longitudinal assessment of 75-year old sample harvested either 3-weeks (**a, b**) or 6-weeks (**c, d**) post-engraftment assessed by immunostaining human-specific protein (**a, c**) and H/E histology (**c, d**). Immunostaining for human-specific spectrin and lamin A/C indicate frequency of central nucleation or peripheral nucleation in regenerated human myofibers 3-weeks (**a**) or 6-weeks (**c**) post-engraftment. Prevalence of interstitial mononuclear cell types indicated at intervals post-engraftment (solid arrows) and their increased presence beyond the boundary of mature human myofibers prevalent in samples 6-weeks post-engraftment (**c, d**; dotted circular region). Within these new regions low-intensity spectrin membrane staining was observed in small-diameter fibers (**c, d**; indicated by ‘+’ signs adjacent to fibers). Asterisks (*) denote muscle orientation between serial cross-sections (**a-d**). **e, f)** Longitudinal assessment of regeneration and nuclear localization after regeneration of the 78-year old sample harvested either 3-weeks (**e**) or 6-weeks (**f**) post-engraftment. Solid arrowheads indicate centrally-nucleated fibers (CNF), while empty arrowheads indicate myonuclei that have adopted a peripheral position within the myofiber (**e, f**). **g, h**) Myonuclei of mouse origin (negative for human lamin A/C) observed within regenerated human spectrin-positive fibers; open arrows identify murine central myonuclei. Scale bars represent 100 μm (**a-f**) or 50 μm (**g, h**).

### Human myonuclei adopt positions subjacent to the sarcolemma following regeneration

In the mouse, central-nucleation provides a clear identifier of regenerated myofibers in mouse skeletal muscle. Interestingly, the myonuclei of regenerated human myofibers quickly adopt a peripheral position subjacent to the sarcolemma (**Figure 6e, f**; *open arrowheads*). This was observed within just 3-weeks following engraftment and had markedly increased in samples harvested 6-weeks post-engraftment, consistent with prior observations ^28^. We previously estimated that some 80% of regenerated mouse muscle fibers contain centrally-positioned nuclei in cross sections ^37^. In regenerated lamin-positive human fibers by contrast, central nucleation represented only a minor fraction of myonuclei marked by human lamin A/C at 6-weeks post-engraftment (**Figure 6f**; *solid arrowheads*). This would either indicate differences in myonuclear domain and spatial distancing in human fibers compared to that in mice or, perhaps, changes in the rates of migration between the central and peripheral positions following myofiber repair or complete regeneration.

### Satellite cells of mouse origin fuse into regenerating human myofibers

One further point of interest arising from microscopic inspection of the border between regenerated human and mouse muscle tissue were rare observations of human myofibers containing central nuclei unmarked by human lamin A/C. While the vast majority of regenerated human myofibers immunostained for both human spectrin and lamin A/C, these ‘mosaic’ myofibers likely resulted from the fusion of mouse with human cells during myogenesis (**Figure 6g, h**). This was seen only in samples collected 6-weeks post-engraftment and suggests that mouse muSCs migrated and fused with actively regenerating human myofibers yielding ‘mosaics’ comprised of both human and mouse myonuclei.

## Discussion

Debilitating effects of waning tissue function with age are currently a central medical issue in the context of aging populations in most ‘developed’ countries, with loss of mass and strength of skeletal muscle in the vanguard. Unsurprisingly therefore, identification of the biological mechanisms behind maintenance of muscle tissue has assumed increasing prominence as a topic of debate and research. Medawar’s seminal essays exploring the fundamental nature of aging ^38,39^, point to a model that predicts an accumulation of functional defects that manifest too late in the life of an individual to significantly affect fecundity and be eliminated from the gene pool. It is to be expected therefore that aging would be a multifactorial consequence of a spectrum of genes operating adversely towards the end of the individual’s lifespan, with few specifically definable culprits. For skeletal muscle, a prominent issue is its loss of mass beyond the 4^th^ decade, centering on the idea of failure of the mechanisms of muscle maintenance, commonly postulated to involve myogenic replacement of muscle lost during day-to-day damage. One level of discussion around this point centers on the extent of such a maintenance function during day-to-day activity. The main evidence on this point, the progressive reduction in telomere length and of *in vitro* proliferative potential in satellite cells, argues strongly against any major implicaton of satellite cell activity beyond adolescence except in myopathic individuals ^23,40^.

Nonetheless, the notion of a functional deficit in the muscle satellite cell population remains a leading candidate to explain age-related sarcopenia, debate centering on whether any demonstrable dysfunction of this cell in the aging muscle is attributable to some intrinsic property of the aging satellite cell itself or on its functional competence in an environment degraded by the ravages of time and subliminal pathology. In the aging mouse, the main experimental model, there is strong evidence for changes in the mechanisms that mediate the activation and restoration of satellite cell dormancy that, between them, would disfavor its role in regeneration. But we lack models in which to establish a direct link of such molecular biological data to a measurable defect in regeneration *in vivo*. Functional defects detected *in vitro* in extracted myogenic cells are conducted in the absence of the plethora of complexities of the *in vivo* situation, while translation of defects in regeneration of mouse muscle are open to doubts as to the relevance to humans due to the fundamental muscle cell biology and inflammatory milieu in the mouse.

In humans, the evidence for an intrinsic defect in aged human satellite cells consists of counts of the numbers of satellite cells present in muscle samples and of their activity *in vitro*. Here, we have attempted to set up the closest situation we could devise to an *in vivo* bout of human muscle regeneration as a means to evaluate the myogenic potential of aged human muscle. As is commonly the case with human research, technical and practical issues obtruded somewhat and we were unable to gather data across as great a range of ages as we had envisaged. It also proved difficult, with several sites of autopsy, to standardize the interval between death and performance of the graft, so we have included this experimental variable in the experimental design. Interestingly, our data shows no obvious effect of postmortem interval on the myogenic performance of the grafts. The implication that the muscle stem cell compartment within the muscle satellite cell population is highly resistant to the anoxic conditions of postmortem muscle accords with a previous demonstration of persistence of myogenic cells in such an environment. This finding resolves an old debate regarding whether the source of myogenic cells in the anoxic center of large muscle grafts is due to survival or by immigration from neighboring less anoxic regions ^41,42^. Clearly, the ability of satellite cells to survive anoxic conditions enhances their value for recovery from large regions of traumatic muscle injury. We were also surprised to have had difficulty obtaining autopsy samples from younger adults, which would have provided a better comparison across the ages. Perhaps this was due to the factors that lead to the decision to perform an autopsy (cause of death).

Nonetheless, our data shows that the very oldest human muscles still contain a population of myogenic cells that were able to reconstruct well-formed muscles with only minimal interstitial fibrosis. This requires well-coordinated choreography of the serial stages of myogenesis in a xenogeneic systemic environment where the inflammatory and vascular systems, as well as cytokine and hormonal signals, differ in detail from those that would operate *in vivo* in man, suggesting that much of the underlying regulation is local to the graft. The near-normal architecture is probably attributable in large part to the persistence of the structural cues of persisting basement membranes. However, we cannot exclude effects of persistent non-myogenic cells, such as fibroadipocytes of human origin and our previous work has shown that some of the microvascular structure is derived from human cells present in the graft ^28^. Whatever the case, our data shows that even aged human muscles remain fully competent to undertake extensive myogenesis in a friendly environment. Taken together with evidence from grafting or parabiosis between animals of different ages ^13,15^, this would implicate the systemic environment, rather than the myogenic cells themselves, or the local interstitial environment within which they undertake regeneration, as the dominant cause of age-related diminution of regenerative response in skeletal muscle.

The model we present here, although imperfect in some respects, provides the closest available simulation of human muscle regeneration *in vivo*. It demonstrates excellent reproduction of normal human muscle architecture within these grafts as a major advantage over grafts of isolated human myogenic cells. The practical and technical problems may be resolved by assembly of a well coordinated supply of cadaver muscle from a single dedicated source, to provide a more stable system with less inter-procedure variability than we encountered. Such a low-noise system woiuld permit investigation of the fine details of human muscle regeneraton that we presently lack which is essential for effective translation of gene and cell therapies from simple mouse models to efficacious human trials.

## Material and Methods

### Study Approval and Human Muscle Samples

Through clinical collaborations and approved Material Transfer Agreements (MTA) with the University of Maryland School of Medicine, Georgetown University Medical Center, Howard University Hospital, and Children’s National Hospital we obtained de-identified human muscle samples harvested during postmortem autopsy procedures and stored in chilled DMEM. Postmortem samples were described only by age, gender, race, and date/time of death and autopsy. Exclusion criteria for human samples included patients with blood-borne infectious diseases (i.e. HIV, Hepatitis) or neuromuscular diseases. This study was approved by the Institutional Review Board (IRB) of Children’s National Hospital and Children’s Research Institute.

### Xenograft Surgical Procedure

Animal procedures were approved by the Institutional Animal Care and Use Committee (IACUC) of Children’s National Hospital and Children’s Research Institute. Female and male NOD.Cg-*Rag1*^*tm1Mom*^ *Il2rg*^*tm1Wjl*^/SzJ (NRG) immunodeficient mice ^35^ aged 2-4 months were used for this study (007799, Jackson Laboratories). Donor human muscle was trimmed into approximately 8 × 4 × 2 mm strips. Mice were anaesthetized with 2-5% isoflurane throughout the procedure. The tibialis anterior (TA) and extensor digitorum longus (EDL) muscles were removed from the anterior tibial compartment, and a strip of human muscle was placed in the empty anterior compartment and ligated with 6/0 POLYPRO non-absorbable sutures (CP Medical) to the tendons of the peroneus longus ^28^. The skin was then closed with Histoacryl surgical glue (B. Braun) and Reflex stainless-steel wound clips (CellPoint Scientific). Buprenorphine SR (1 mg/kg) was administered subcutaneously post-surgery for pain control.

### Histology and Immunostaining

Frozen muscles were sectioned at 8 μm thicknesses using a Leica CM1950 cryostat maintained at −20°C. The muscle sections were first stained with hematoxylin and eosin (H/E) for histological assessment of xenograft regeneration. Muscle sections were also immunostained with human-specific spectrin (SPEC1-CE, 1:50, Leica) and human-specific lamin A/C (ab40567, 1:200, Abcam), while total muscle tissue was marked by 4’6-diamidino-2-phenylindole (DAPI) staining (outside human-marked tissue) or by immunostaining with fluorescently-conjugated wheat germ agglutinin (WGA alexafluor-647, W32466, 1:500, Thermo Fisher Scientific). Specifically, muscle sections were fixed in ice-cold methanol for 10 minutes, washed in Phosphate-Buffered Saline (PBS), and blocked in PBS supplemented with 2% goat serum (GTX73249, GeneTex) and mouse-on-mouse (M.O.M.) blocking reagent (MKB-2213, Vector Laboratories). Muscle sections were incubated with primary antibodies overnight at 4°C and subsequently probed with Alexa Fluor secondary antibodies (Life Technologies). Finally, muscle sections were mounted with Prolong Gold Mounting Media (Life Technologies) with DAPI for nuclear staining.

### Muscle Fiber Analyses

Human regenerated skeletal muscle fibers were quantified in ImageJ software and defined by human-specific spectrin membrane staining and human-specific lamin A/C nuclear staining. Central or peripheral myonucleation was assessed based on the position of myonuclei labeled with human-specific lamin A/C. Minimal Feret diameter was used to calculate muscle fiber size due to its strength in limiting experimental error that may arise from tissue orientation or the sectioning angle ^43^.

### Microscopy

Images were acquired on the Olympus BX61 VS120-S5 Virtual Slide Scanning System Microscope with UPlanSApo 20x/0.75 and 40x/0.95 objectives, Olympus XM10 monochrome camera, and Olympus VS-ASW FL 2.7 imaging software. Analysis was performed using Olympus CellSens 1.13 and ImageJ software.

## Data Availability

The authors declare that the main data supporting the findings of this study are available within the article and its Supplementary Information files. Additional data are available from the corresponding author upon request.

## Acknowledgements

We would like to thank The University of Maryland School of Medicine Pathology Department, The Georgetown University Medical Center Pathology Department, The Howard University Hospital Pathology Department, and the Children’s National Hospital Pathology Department for providing the human muscle samples utilized in this study. Funding for this project was obtained by TAP and EPH. This work was supported by the National Institutes of Health NIAMS (R21AG051260 | TAP, EPH; T32AR056993 | JSN, DAGM), Foundation to Eradicate Duchenne (JSN, DAGM), Muscular Dystrophy Association (MDA295203 | TAP; MDA480160 | JSN), Parent Project Muscular Dystrophy (TAP), and Duchenne Parent Project Netherlands (JSN). Microscopy imaging was performed at the Children’s National Hospital/Children’s Research Institute Cellular Imaging Core, which is supported by funds from Children’s Research Institute and The National Institutes of Health NICHD (U54HD090257).

## Authorship Contributions

This study was initially conceptualized by TAP, EPH and KRW. Xenograft procedures were performed by JSN, DAGM, MN and TAP in accordance with Institutional IRB and IACUC approvals acquired by MN, JSN and TAP. Human muscle samples were provided through collaborations with OBI, BTH, MNFL, CR and DAH, together with their respective clinical pathology departments. Cellular analyses, including histology and immunostaining, were performed by JSN, DAGM, MN, NFH and TD. The manuscript was written by JSN, DAGM, and TAP and reviewed by all contributing authors. JSN and DAGM share co-authorship on this manuscript, having contributed significantly to the execution, analysis and publication of this study. The order of first authorship was arranged according to contributions made to study development and execution, data analysis and publication of the results. JSN and TAP share co-corresponding authors on this manuscript.

